# Single cell transcriptomics reveals enrichment of aggregation-prone alpha-synuclein isoforms across synucleinopathies

**DOI:** 10.1101/2025.10.13.682119

**Authors:** E. Keats Shwab, Webb Pierson, Daniel C. Gingerich, Zhaohui Man, Sapir Margalit, Dean Yona, Asaf Sivan, Julia Gamache, Geidy E. Serrano, Thomas G. Beach, Yuval Ebenstein, Roy Beck, Ornit Chiba-Falek

**Author notes:** These authors contributed equally to this study (E. Keats Shwab, Webb Pierson). To whom correspondence should be addressed: Ornit Chiba-Falek Division of Translational Brain Sciences, Dept of Neurology Duke University School of Medicine Durham, North Carolina 27710, USA.

## Abstract

Alpha-synuclein (α-Syn) is the primary component of Lewy bodies, the pathological hallmark of neurodegenerative synucleinopathies, including Parkinson’s disease (PD) and dementia with Lewy bodies (DLB). Dysregulated expression of its encoding gene, *SNCA*, has been identified in association with both PD and DLB in short-read sequencing studies. However, such studies do not capture variation in transcript isoforms expressed. Here we combine for the first time *SNCA*-targeted long-read multiplexed arrays isoform sequencing (MAS-Iso-seq) with unbiased short-read single nucleus (sn) RNA-seq for simultaneous characterization of the *SNCA* transcript isoform landscape and mapping of isoform expression to specific cell types and subtypes in PD, DLB, and control sample cortical tissues. This approach enabled discovery of numerous *SNCA* transcript isoforms displaying novel splicing patterns and incorporating novel exons. We further identified an abundant class of transcript isoforms encoding a previously unreported α-Syn protein variant (α-Syn-115) and displaying increased proportional detection in excitatory neurons of PD and DLB tissues in comparison to controls. The proportion of these isoforms was found to be especially high within several specific glutamatergic neuron subtypes. In-depth characterization of the predicted structural and biochemical properties of α-Syn-115 using an *in silico* modeling approach revealed a greater aggregative affinity compared with canonical α-Syn-140, suggesting the potential for increased cytosolic α-Syn-115 abundance to induce aggregation between heterogeneous α-Syn isoforms, potentially driving fibril formation and disease progression. Together, our findings provide new insights into the molecular mechanisms underlying α-Syn involvement in multiple synucleinopathies and have translational implications for the development of new precision medicine strategies to combat these diseases, indicating the potential for treatments targeting both specific transcript and protein isoforms, as well as disease-driving cell subtypes.

## INTRODUCTION

Alpha-synuclein (α-Syn) protein is the primary component of Lewy bodies and Lewy neurites, the pathological hallmarks of synucleinopathies including Parkinson’s disease (PD) and dementia with Lewy bodies (DLB)^1^. Moreover, multiple lines of evidence have implicated its coding gene, *SNCA,* as a strong genetic factor in both PD and DLB: autosomal dominant mutations within the *SNCA* coding region were discovered in the rare familial form of PD (fPD)^2^, and genome-wide association studies (GWAS) have demonstrated high correlation of *SNCA* loci with the common non-Mendelian forms of PD^3,4^ and DLB^5^. Analyses of the human brain transcriptome using bulk and single-cell approaches have suggested dysregulation of *SNCA* gene expression as an underpinning disease mechanism^6–9^, which has been supported by studies using disease model systems^10,11^. However, such studies are only able to provide information about gene expression levels and do not capture variation in transcript isoforms expressed and hence the landscape of the various protein products.

Transcriptional and post transcriptional mechanisms, including alternative exon splicing, alternative transcriptional start sites (TSS), and alternative polyadenylation sites may give rise to a variety of proteins translated from a single gene. While changes to 5’ and 3’ untranslated regions (UTR) may affect the level of the protein via RNA stability and translation efficiency, alternative splicing may result in divergent protein isoforms exhibiting truncations, extensions, and other changes to amino acid composition. Noteworthily, alternative transcript splicing has been previously linked to PD and other neurodegenerative disorders such as Alzheimer’s disease (AD), amyotrophic lateral sclerosis (ALS), frontotemporal dementia, and repeat expansion diseases^12,13^.

The *SNCA* gene comprises six canonical exons, of which exons 2-6 are protein encoding, with exon 1 consisting of 5’-UTR. The gene region includes a large intron of > 90kb between exons 4 and 5 (intron 4). The most abundant α-Syn protein isoform (α-Syn-140) is 140 amino acids (aa) in length^14,15^ and is encoded by splicing of all six exons. The full length α-Syn-140 is present in neurons as structured membrane-bound and intrinsically disordered cytosolic forms^16^, and undergoes a range of conformational changes based on the specific architecture of the membranes with which it associates^14,17,18^. When membrane bound, the structurally rigid N-terminus (roughly residues 1-25, encoded from exon 2) constitutes a membrane anchoring region, while the central residues ∼26-100 (exons 2-4) display dynamic structural properties and are involved in sensing of membrane architecture^14,19^. Missense mutations changing the residues in this region have been implicated in aggregation and autosomal dominant fPD^20–22^. The C-terminal residues ∼100-140 (exons 5-6) are highly disordered and likely exhibit only transient membrane interaction^14,19^. Negatively charged residues within this C-terminal region have been demonstrated to counteract aggregation^23^.

At least three additional alternatively spliced transcripts were previously described in humans^24^. Skipping of exons 3, 5, or both results in in-frame deletions, and alternative transcripts encoding proteins of 126^25^, 112^25–28^, and 98^29,30^ aa, respectively. The protein products resulting from these alternative splicing events acquire different properties. Isoforms with truncated C-termini, such as α-Syn-112 and -98, show an increased aggregation rate^23^. Conversely, as α-Syn-126 lacks a portion of the central region involved in fibril formation but retains the negatively charged C-terminus, this isoform is predicted to exhibit a reduced aggregative propensity^23^. A protective role for the shortened α-Syn-126 is supported by reduced expression of this transcript isoform in frontal cortices of DLB patients, whereas transcripts for the aggregation-prone isoforms, α-Syn-112 and -98, were found to be upregulated in these same samples^31^. In addition, three alternative 5’ noncoding exons (2a, 2b, and 2c) have been described as alternative 5’-UTRs to exon 1^32^.

Previously, we catalogued *SNCA* transcript isoforms expressed in bulk temporal cortex (TC) samples of PD, DLB and normal control (NC) donors using long-read sequencing with targeted capture of *SNCA* cDNA^7^. In that study, we identified a total of 41 known and novel unique transcript isoforms. Transcripts translating to known α-Syn isoforms included 28 predicted to encode α-Syn-140 with variable 5’-and/or 3’-UTR sequences, 2 encoding α-Syn-126, and 5 encoding α-Syn -112. In addition, our analysis identified 2 isoforms predicted to encode a novel 115aa protein (α-Syn-115) due to the introduction of an alternative stop codon located in intron 4. However, the primary limitation of this bulk sequencing methodology was that we were unable to identify the cell types expressing each specific isoform. To gain mechanistic insights about the roles of different *SNCA* transcripts in disease, it is imperative to comprehensively characterize the complete landscape of *SNCA* transcripts including the less abundant isoforms, to identify the cell types expressing each specific isoform, and to examine the relative abundance of transcript expression within cell types and across disease states.

Here, we address these gaps and for the first time characterize the entire full-length transcriptomic landscape of the *SNCA* gene at a single-cell resolution. To accomplish this, we integrated two cutting-edge transcriptomic technologies: (1) long-read multiplexed arrays isoform sequencing (MAS-Iso-seq)^33^, enriched for *SNCA* transcripts to identify all transcript isoforms including rare transcripts, and (2) short-read single-nucleus (sn) RNA-seq^34^ for cell subtype annotation and quantitation of gene expression levels to determine the cell types in which each transcript isoform is expressed. We used the same barcoded cDNA preparations for both sequencing libraries, allowing us to integrate the short-and long-read sequencing datasets. This approach enabled identification of specific *SNCA* isoform expression in individual nuclei annotated not only at the major cell type level but also at the subtype level. The outcomes provided an unprecedented insight into the molecular mechanisms underlying synucleinopathies as well as critical knowledge for *SNCA*-targeted drug development to fight these important diseases.

## RESULTS

### Characterization of cell types and subtypes in the human temporal cortex (TC) of individuals with PD, DLB, and controls using short-read sequencing datasets

After quality control (QC) filtering, short-read snRNA-seq datasets from 98,046 NC nuclei, 109,932 PD nuclei, and 103,906 DLB nuclei were retained from TC brain samples of 12 donors for each disease state (**Tables S1** and **S2** summarize the demographic and neuropathological phenotypes of donor samples). Gene expression data from NC nuclei were separately integrated with PD and DLB nuclei, then annotated according to major brain cell types by label transfer^35^ from a pre-annotated reference snRNA-seq dataset^36^ **(Fig. 1A)**. Among all NC, PD, and DLB nuclei together, annotated nuclei included 7.50% astrocytes, 23.51% excitatory neurons, 10.15% inhibitory neurons, 6.38% microglia, 47.41% oligodendrocytes, and 5.03% oligodendrocyte precursor cells (OPCs). Other cell types, including endothelial cells and vascular and leptomeningeal cells, made up less than 1% of the total cell population and were therefore excluded from the dataset in downstream analyses.

**Figure 1.**
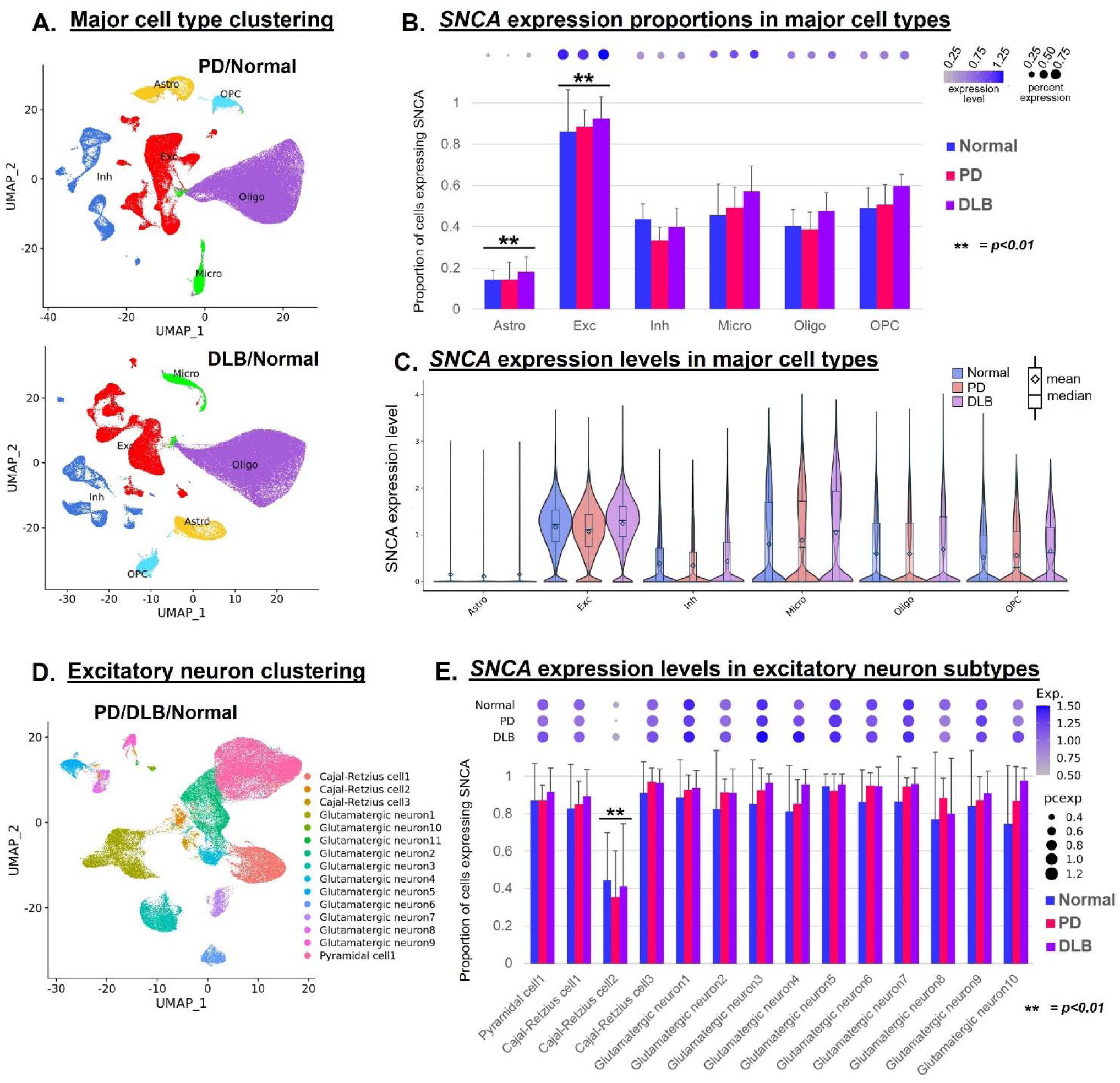
Patterns of *SNCA* expression in Normal, PD, and DLB cell types and subtypes. **A.** UMAP dimensional reduction plots of snRNA-seq transcriptomic data for PD (upper panel) and DLB (lower panel) nuclei integrated with NC nuclei. Plots are color coded to indicate nuclei annotated as astrocytes (Astro), excitatory neurons (Exc), inhibitory neurons (Inh), microglia (Micro), oligodendrocytes (Oligo) and oligodendrocyte precursor cells (OPC). **B.** Histogram showing proportions of nuclei of each major cell type and disease state expressing *SNCA*. Bars represent mean proportions of expressing nuclei for each donor sample within corresponding disease state/cell type. Error bars represent standard deviations. **= p<0.01, Comparisons of pooled diagnosis groups between the 6 cell types via ANOVA with post-hoc paired Tukey HSD tests. Dot plots above histograms indicate SNCA expression level (color saturation) and proportion (dot size). **C.** Violin plots showing SNCA expression levels for individual nuclei within each disease state/cell type. Box plots represent upper and lower quartiles, with center lines indicating data median and diamonds indicating means and whiskers indicating data spread. **D.** UMAP dimensional reduction plot of snRNA-seq transcriptomic data for excitatory neurons of PD and DLB nuclei integrated with NC nuclei. Plots are color coded to indicate subtype clusters with MayoMap annotation. **E.** Histogram showing proportions of nuclei of each excitatory neuron subtype and disease state expressing *SNCA*. Features are as in panel B.

In order to explore the regulation of *SNCA* expression in relation to synucleinopathies, we first determined which major cell types were predominantly expressing the *SNCA* gene across disease states. We examined the proportion of PD, DLB and NC nuclei exhibiting *SNCA* expression counts > 0 within each cell type. Excitatory neurons were found to express *SNCA* in a significantly higher proportion of cells compared to each of the other cell types (*p* < 0.01), with mean expression of approximately 80-100% per sample, while astrocytes showed expression in the significantly lowest proportion of cells compared to other cell types (*p* < 0.01), approximately 20% on average, and expression proportion in other cell types ranged between 40 to 60% **(Fig. 1B**, **Table S3)**. Consistently, the highest mean and median *SNCA* expression levels were found in excitatory neurons, with the second highest in microglia and the lowest in astrocytes **(Fig. 1B)**. We didn’t find significant differences in *SNCA* expression proportions between the NC, PD, and DLB diagnosis groups for any of the major cell types (*p* > 0.05) **(Table S3)**. We also examined *SNCA* expression at a cell subtype level by delineation of subtype clusters within integrated PD/NC and DLB/NC nuclei datasets **(Fig. S1A)**. Across the 35 and 32 clusters of the respective PD/NC and DLB/NC integrated nuclei, *SNCA* expression proportions and levels were generally higher within excitatory neuron clusters than other cell subtypes **(Fig. S1B)**.

Next, we focused on patterns of *SNCA* expression in neurons only and directly compared the same neuronal subtype populations between NC, PD, and DLB nuclei. Data integration and clustering of excitatory neuron nuclei only from all three pathologies, NC, PD, and DL,B identified 14 subtypes, annotated by scMayoMap^37^ **(Fig. 1D)**. Analysis revealed relatively uniform proportions and levels of *SNCA* expression across the excitatory neuron subtypes, with significantly lower *SNCA* expression proportion only in Cajal-Retzius cell 2 subtype compared to each of the other subtypes (*p* < 0.01) **(Fig. 1E**, **Table S4)**. A similar analysis was carried out for inhibitory neuron nuclei; amongst 23 subtype populations **(Fig. S1C)**, *SNCA* expression proportions and levels were more varied **(Fig. S1D)**.

*SNCA-AS1*, the antisense transcript to the *SNCA* gene, has been previously shown to play an important role in aging and PD, in part through regulation of the expression of *SNCA* and numerous other genes^38^. For this reason, we examined its patterns of expression among major cell types within our snRNA-seq dataset. As for *SNCA*, we found that a significantly higher proportion of excitatory neuron cells expressed *SNCA-AS1* compared to the other major cell types across the disease states **(Fig. S2A,B, Table S5)**. Moreover, expression percentage was qualitatively increased in PD compared to NC and DLB excitatory neurons, but this difference was not found to be statistically significant. Overall, only a small proportion (< 1.5%) of cells of any type were found to express *SNCA-AS1* on average across donor samples. When expression proportions within specific excitatory neuron subtypes were examined, mean expression proportion was consistently higher in PD compared to both NC and DLB cells in ten of 13 subtypes **(Fig. S2C,D)**.

### Identification of *SNCA* transcript isoforms expressed in PD, DLB and NC brain nuclei using long-read MAS-Iso-seq

Transcriptional and post-transcriptional processes regulate gene expression levels but also give rise to various transcript isoforms^39^. While our short-read snRNA-seq data provide valuable quantitative information on expression levels of *SNCA*, this is limited to the gene level, without differentiating between *SNCA* transcript isoforms. We were next interested in characterizing the full range of specific *SNCA* isoforms expressed. Therefore we employed long-read sequencing, using MAS-Iso-seq technology^33^, in conjunction with targeted enrichment of *SNCA*-mRNAs. A subset of the same cDNA samples processed for the short RNA-seq experiments were used to prepare the long-reads snRNA-seq libraries, thus enabling direct correlation between long-read and short-read sequencing data. We performed MAS-Iso-seq using 11 of the same cDNA libraries used above for snRNA-seq, including four each from TC samples of NC and DLB donors, and 3 from PD donors, as well as a cDNA library from 1 additional PD donor not used for snRNA-seq, for a total of 4 samples for each diagnosis group **(Table S1)**. Following processing and QC filtering of MAS-Iso-seq reads, we identified 327 total unique *SNCA* transcript isoforms expressed among the 12 donor samples. We then grouped the identified 327 unique transcripts into 62 isoform classes based on RNA processing and coding attributes: (1) exons included, (2) 5’ and 3’ splice sites for individual exons, and (3) the presence of open reading frames (ORFs) predicted to encode structured α-Syn (based on N-terminal membrane-binding domain) **(Fig. 2A)**. Isoform classes were assigned a unique letter (A-Z) based on exon inclusion, and subclasses based on specific exon splice sites were delineated by numbers (*e.g.* A.1 and A.2). Of note, cDNA molecules were generated by 3’ to 5’ extension via reverse transcription from polyA tail-annealing primers and thus may have incomplete 5’ ends. Isoform subclasses were therefore annotated as potentially incomplete (denoted by lowercase “i”) unless they included 5’ UTR upstream of the canonical translation start codon in exon 2 (*e.g.* D.1i vs. D.1). Lowercase letters “*a*-*d*” were used to further distinguish multiple incomplete transcripts matched to the same complete transcript (*e.g.* A.1i*a* and A.1i*b*), and transcripts including ORFs encoding structured α-Syn (*e.g.* B.1*a* vs. B.1*b*).

**Figure 2.**
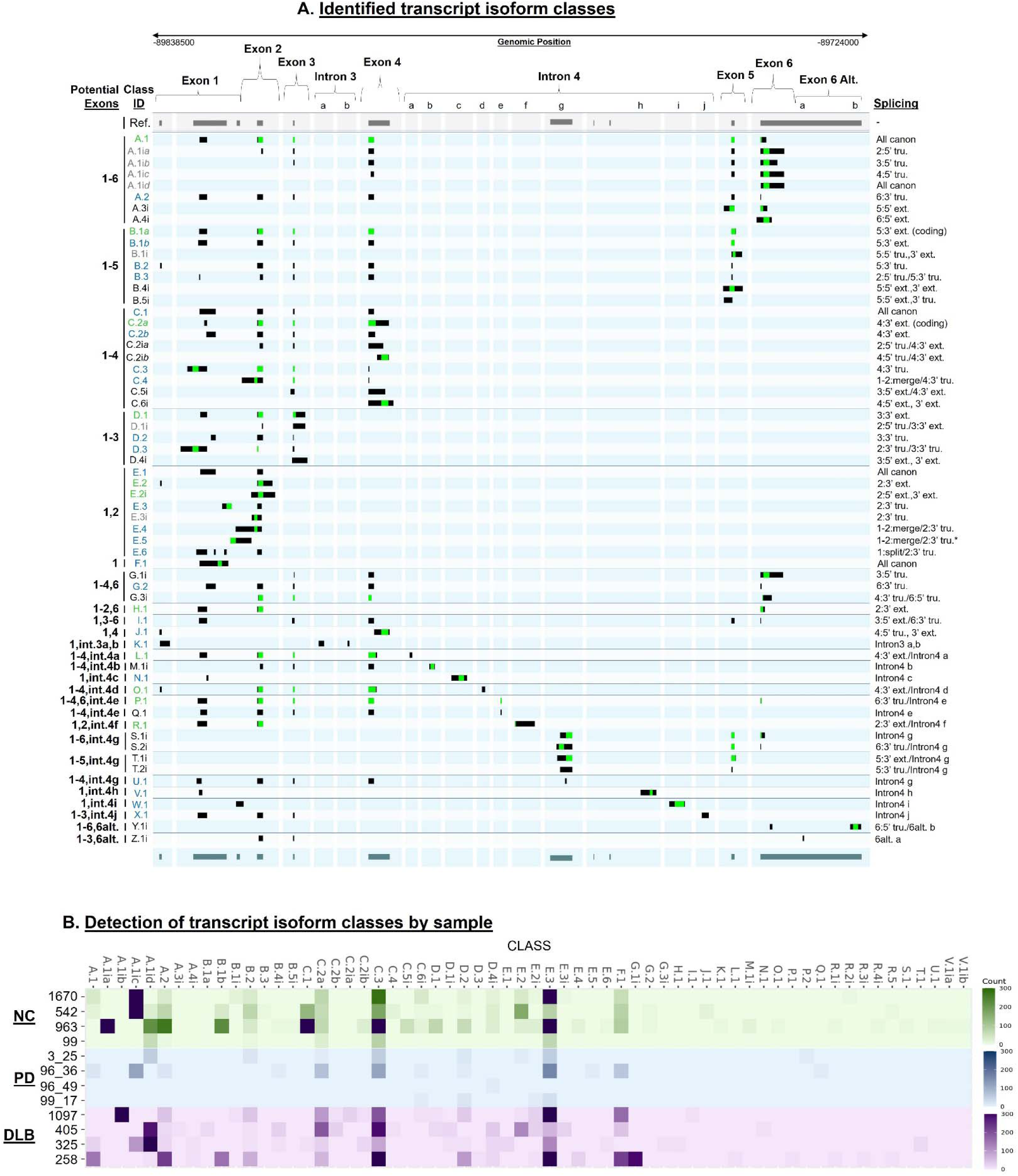
Identified transcript isoform splicing classes. **A.** Genomic track plots showing exon regions for a representative transcript from each splicing category. Top arrow indicates chromosome 4 region covered by track plots. Exon and intron regions are indicated by brackets. Lowercase letters within intron 3 and 4 regions indicate novel exons. Grayscale bars at top represent previously reported (reference) exonic regions. Alternating blue and light blue rows show unique isoform splicing categories, with identifiers shown to the left side. Lowercase ‘i’ in IDs indicates potentially incomplete isoforms. Green IDs indicate isoform categories encoding SNCA protein with exon 2-encoded membrane-binding domain. Blue IDs indicate complete isoform categories not encoding protein with membrane-binding domain. Black IDs indicate unique potentially incomplete isoform categories (lacking 5’ UTR). Gray IDs indicate potentially incomplete isoform categories that may represent incomplete versions of complete categories with the same ID. Lowercase italicized ‘*a-d*’ indicate different incomplete transcripts that align to the same complete transcript. Exon numbers included in isoform categories are shown to the far left of the figure. Specific splicing attributes are shown to the far right. **B.** Heatmap showing log-normalized counts per donor sample of transcripts from each isoform splicing category shown in panel A, excluding noncoding mono-exonic isoforms. NC samples are shown in shades of green, PD samples in shades of blue, and DLB samples in shades of purple.

Overall, we identified a wide variety of *SNCA* transcripts **(Fig. 2A)**, including those comprising the full set of 6 canonical exons (class A), as well as 9 additional combinations of canonical exons (classes B-J). These latter included classes of isoforms with truncated 3’ ends (B-F), ranging from loss of exon 6 alone (B) to the loss of all exons except exon 1 (F). We also identified several classes of isoforms exhibiting skipping of particular canonical exons (G-J). This included skipping of exon 5 (G), exons 3-5 (H), exons 2 and 5 (I), and exons 2, 3, 5 and 6 (J). Interestingly, we did not identify any isoforms skipping only exon 3, or only 3 and 5 together, as have been previously reported^25,29,30^. We did identify numerous isoforms that incorporated noncanonical exons (classes K-Z), including 2 novel exons within the intron 3 region (K), and 10 distinct novel exons within the large intron 4 region (J-X), as well as 2 novel exons 3’ to canonical exon 6 (Y, Z). A high degree of variability was observed in the exon 1 5’ UTR, which was not taken into account in isoform classification (to avoid excessive complexity and due to TSS uncertainty resulting from the aforementioned 3’ bias in cDNA generation) except in cases where exons 1 and 2 were merged (*e.g.* E.4), or where exon 1 was split into multiple exons (E.6). 3’ UTR variability was also observed, mainly stemming from variable extension of the exon 6 3’ end, but also from the inclusion of novel alternative downstream exons in classes Y and Z.

In comparison to our previous study^7^ using bulk tissue MAS-Iso-seq, we identified an overall greater diversity of isoforms in the current study, possibly reflecting greater sensitivity of the current methodology to detect less abundant transcripts. Of the combinations of canonical exons detected in this study described above, only classes A (all canonical exons) and G (exon 5 skipping) were previously represented. However, in the previous study, isoforms skipping only exon 3, or skipping both 3 and 4 together were identified, which were not detected in the current study. Of the novel exons detected in the current work, only novel exon g within intron 4 was identified previously, although an additional intron 4 exon was identified previously that is not found in the current dataset. No alternative exons within the intron 3 region or downstream of exon 6 were previously detected. Additionally, in the current work we detected greater variability of both 5’ and 3’ splice sites for each exon than were previously identified, with only alternative splicing of exons 1, 2 and 4 described in the previous study, whereas in the current work we identified alternative splicing of all canonical exons, with both 5’ and 3’ extension and truncation observed for most. In both studies, isoforms with alternative 3’ ends due to 3’ extension of exon 4 and loss of exons 5 and 6 were identified (class C.2*a* in the current study).

Quantitative analysis, by sequencing read count measurement, showed that E.3 was the most consistently detected sub-category across all donor samples from all pathological groups, followed by C.2*a* and C.3 **(Fig. 2B**, **Table S6)**. Isoform A.1, comprising the full set of canonically spliced exons, was only detected in 2 NC, 2 DLB, and 1 PD sample. However, subclasses A.1i*a-d* may represent incomplete sequences of A.1 and together were identified more strongly and across a wider sample set. The other isoforms in the dataset had varying levels of detection, with no single isoform detected across all samples, and many isoforms detected in only one sample each, particularly those isoforms that include novel exons, suggesting low expression and/or rapid mRNA degradation.

To examine variation in *SNCA-AS1* transcripts across disease states, we analyzed our MAS-Iso-seq data to catalog the different *SNCA-AS1* transcript isoforms expressed. In doing so, we identified six different isoforms, which we termed *SNCA-AS1* A-F **(Fig. S2E, Table S7)**. Isoforms A-E comprised partially overlapping genomic regions with one another, while isoform F overlapped only with B and E. Each *SNCA-AS1* isoform was detected in only one donor sample, and only NC sample 963 showed expression of more than one isoform (A and B) **(Fig. S2F)**. Isoforms A-C were identified in NC cells, isoform D in PD cells, and isoforms E and F in DLB cells.

### Prediction of known and novel a-Syn proteins encoded by the full-length *SNCA* transcript isoforms

Of the 62 classes of transcripts isoforms defined via the long-read analysis described above, 11 were found to contain ORFs predicted to encode α-Syn protein with the structured N-terminal domain encoded on exon 2, responsible for membrane binding and thus required for normal α-Syn functionality. These 11 classes were together predicted to encode 8 distinct α-Syn proteins of varying aa lengths **(Fig. 3A)**. These include the canonical α-Syn-140 (encoded by class A.1), as well as C-terminally truncated species ranging from 138 to 42aa in length. Class B.1*a* encodes a 138aa protein (α-Syn-138) resulting from 3’ extension of exon 5, generating a premature stop codon. Class P.1 encodes α-Syn-120, resulting from replacement of exon 5 with a novel exon 4e within the intron 4 region, which also causes a frameshift in the exon 6 coding region with an alternative stop codon. Notably, a 115aa protein (α-Syn-115) is encoded by 3 different isoform classes, C.2*a*, L.1, and O.1, with 3’ extension of exon 4 resulting in a premature stop codon and an extended 3’ UTR, which could have a stabilizing influence for the mRNA molecule. This may partially explain why class C.2*a* isoforms were among the most consistently detected across donor samples **(Fig. 2B)**. Classes L.1 and O.1 also include splicing of novel noncoding exons 4a and 4d within the intron 4 region, contributing additional alternate 3’-UTR to the transcript. α-Syn-115 would potentially have a similar structure and properties to the previously reported α-Syn-112, as both lack the exon 5-encoded negatively-charged C-terminus which helps protect against aggregation^23^, but retain the central region encoded on exons 3 and 4 associated with increased aggregation^20–22^. Thus, α-Syn-115 may have a similar increased tendency towards fibril formation as has been demonstrated for α-Syn-112^23^. Additional predicted proteins include: 2 distinct 64aa species (α-Syn-64A and -64B), resulting from 3’ extension of exon 3 (D.1) and skipping of exons 3-5 (H.1), respectively; a 48aa protein (α-Syn-48) resulting from inclusion of a novel exon 4f in the intron 4 region, and introduction of an alternate stop codon and long 3’ UTR (R.1); and a 42aa protein (α-Syn-42) resulting from 3’ exon 2 extension in classes E.2 and E.2i. In addition to the isoform classes mentioned above, some incomplete classes may also encode these or other α-Syn proteins. For example, protein α-Syn-140 is predicted to be encoded by isoform category A.1, but may also be encoded by the complete forms of incomplete transcripts A.1i*a-d*. Likewise, while no isoform classes were identified encoding α-Syn-112, class G.1i may represent an incomplete sequence of an α-Syn-112-encoding transcript, as exon 5 is skipped while the remainder of the present sequence shows a canonical splicing pattern downstream of a 5’ truncated (possibly due to incomplete cDNA generation) exon 3. However, no incomplete isoforms identified here are likely to encode the previously described α-Syn-126 or α-Syn-98 proteins.

**Figure 3.**
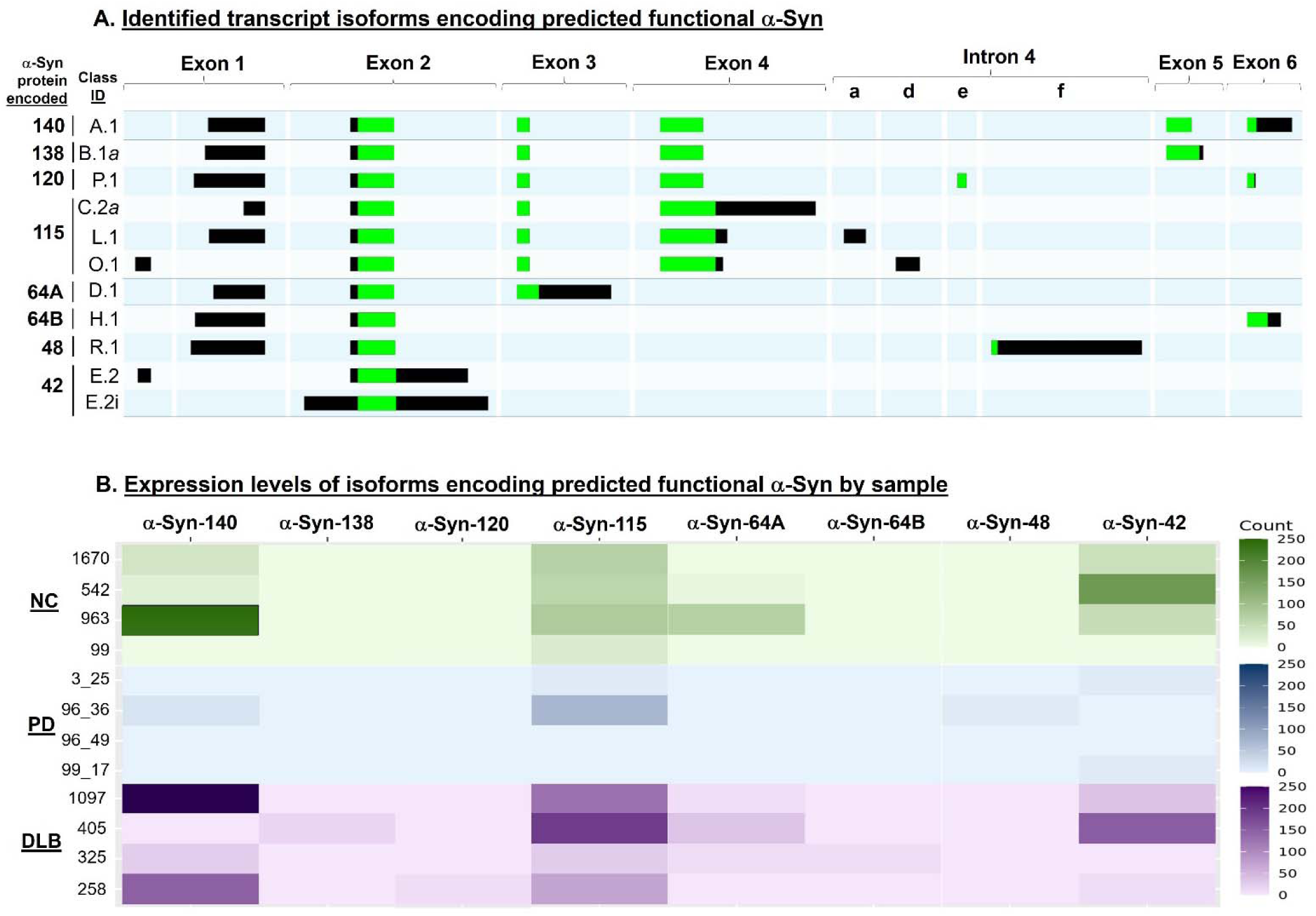
Transcript isoform splicing categories predicted to encode functional a-Syn protein. **A.** Genomic track plots showing exon regions for a representative transcript from splicing categories predicted to encode α-Syn protein with exon 2-encoded membrane-binding domain. Exon and intron regions are indicated by brackets. Lowercase letters within intron 4 region indicate novel exons. Alternating blue and light blue rows show unique isoform splicing categories, with identifiers shown to the left side. Predicted encoded protein lengths are shown to the far left of the figure. **B.** Heatmap showing log-normalized counts per donor sample of transcripts from each isoform splicing category predicted to encode the α-Syn protein variant indicated. This includes isoforms of categories A.1, A.1i*a*, A.1i*b*, A.1i*c* for α-Syn-140, B.1*a* for α-Syn -138, P.1 for α-Syn -120, C.2a, C.2i*a*, L.1, and O.1 for α-Syn -115, D.1 for α-Syn -64A, H.1 for α-Syn -64B, R.1 for α-Syn -48, E.2 and E.2i for α-Syn -42. NC samples are shown in shades of green, PD samples in shades of blue, and DLB samples in shades of purple.

Comparison of the detected abundance of protein-encoding transcript isoforms among donor samples and disease states is shown in **Fig. 3B** and **Table S6**. Notably, α-Syn-115-encoding isoforms were detected in the highest number of samples, followed by α-Syn-42. Α-Syn-140 encoding isoforms were detected in fewer samples, but had comparatively high levels of detection in several samples. It should also be noted that shorter transcripts may be artificially more abundant due to the potential for incomplete cDNA generation of longer transcripts, so accurate comparison of abundance of different isoforms is difficult. However, these results do suggest consistent expression of α-Syn-115 and α-Syn-42 encoding isoforms across samples and disease states.

### Structural modeling of predicted a-Syn protein variants

To follow up on the functional implications of the 8 predicted protein variants discussed above in disease context, we investigated the proteins’ structure using an *in silico* structural modelling approach and specifically analyzed the membrane-bound features of each predicted isoform **(Fig. 4A)**. All 8 predicted α-Syn proteins share a common N-terminal region, which consistently adopted an α-helical structure as in the membrane-bound native form. However, the overall per-residue model confidence (pLDDT) was low across many regions **(Fig. S3A)**, consistent with the classification of a-Syn as an intrinsically disordered protein (IDP). The shared predicted α-helical structures in the N-terminal region of all isoforms modeled the known membrane-binding state; however, these models do not fully capture the conformational diversity expected of intrinsically disordered proteins in solution, and therefore should be interpreted cautiously. Notably, the α-Syn-115 predicted structure included additional helical regions not observed in α-Syn-140 **(Fig. 4A)**. However, these regions also exhibited low confidence scores **(Fig. S3A)**. Complementary predictions of the level of structural disorder^40^ suggested that in α-Syn-115, these regions are more structured than in the canonical isoform **(Fig. S3B)**, supporting the potential presence of additional secondary structure elements.

**Figure 4.**
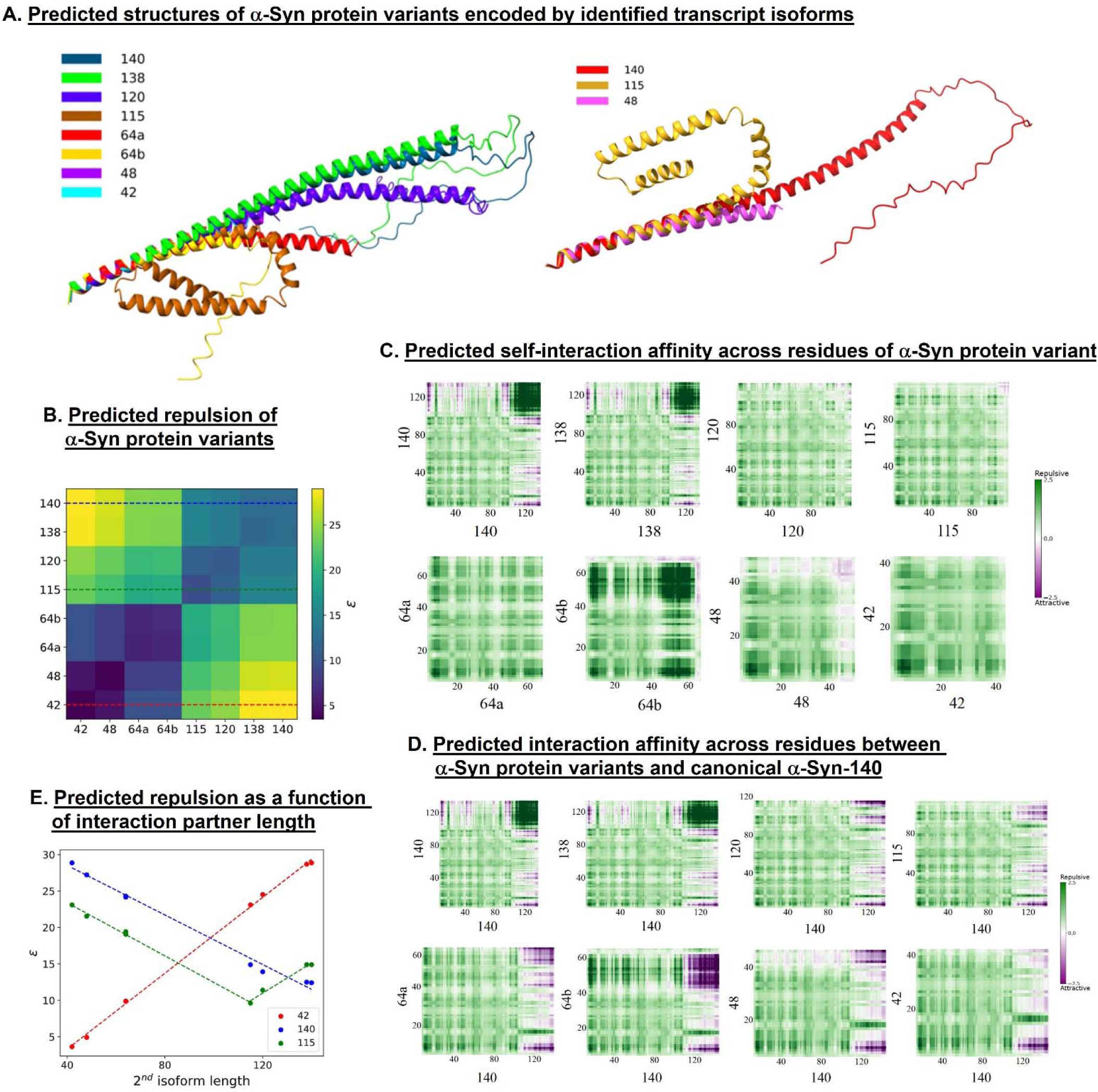
Structural modeling of predicted a-Syn protein variants. **A.** In-silico membrane-bound structural modeling the a-Syn protein isoforms. The N-terminal membrane binding domain is preserved throughout the isoforms, but the 115 long isoforms show additional helicity in the C-terminal domain which isn’t present in other isoforms. Longer isoforms have more disordered ends, while shorter ones are completely ordered. **B.** Mean-field intermolecular interaction parameter estimation (ε) between two isoforms. Near-linear increase in self-repulsion is shown with length (see also Fig S4). The dashed lines signify which ε values are shown in panel E. **C.** Predicted intermolecular maps between identical isoforms. Isoforms 140 and 138 have two sections that are repulsive to themselves, but attractive to one another. The other isoforms are also generally repulsive but with attractive domains. **D.** Predicted intermolecular map between different isoforms and a-Syn-140. All isoforms have attractive sections with the canonical a-Syn-140 isoform. **E.** Estimated repulsion of a-Syn-42 (red), a-Syn-115 (green), and a-Syn-140 (blue) with other isoforms. The values of ε between two different isoforms scale linearly with the partner’s length. For shorter/longer isoform partner, ε decreases/increases linearly.

To further interpret the predicted structures and explore how sequence properties may influence molecular interactions, we analyzed the physiochemical properties of each isoform from its amino acid sequence. Hydropathy profiles revealed that α-Syn-115 displays increased local hydrophobicity at the C-terminus **(Fig. S3C)**. The α-Syn-140 sequence is characterized by a positively charged N-terminal region and a negatively charged C-terminal region and a potential intramolecular electrostatic attraction between termini^41^. In contrast, charge profiles indicated that all of the alternative proteins with the exception of α-Syn-138 lack the negatively charged C-terminal tail, which may significantly alter their interaction dynamics and aggregation potential **(Fig. S3D)**.

As fibril formation occurs cytosolically, we further characterized intermolecular interactions for the 8 proteins in the membrane-unbound and structurally disordered state, revealing significant differences between the proteins. We used the FINCHES algorithm^42^ to quantify the overall attractive or repulsive tendency (ε) between two protein sequences. All interactions between a-Syn isoforms resulted in positive ε values, ranging from weak to strong repulsion **(Fig. 4B)**. However, α-Syn-115 exhibited weaker repulsion than α-Syn-140 or -138, suggesting a potentially increased tendency for self-association or altered interaction behavior compared to the canonical isoform. Additionally, while C-terminal repulsion is predicted in self-interactions of a-Syn-140 and -138, in shorter isoforms (excepting α-Syn-64B), this repulsion is markedly reduced **(Fig. 4C)**, and in cross interactions between a-Syn variant proteins and α-Syn-140, attraction is predicted between shorter isoforms and the C-terminus of the full length α-Syn-140 **(Fig. 4D)**. Such local attraction could potentially induce aggregation between nearby isoforms in heterogeneous solution.

Furthermore, we found that mean-field self-interaction parameter estimations scale somewhat linearly with isoform length **(Fig S3E)**. Moreover, for a given isoform, the repulsion it experiences from different isoforms also scales linearly with the isoform partner length **(Fig. 4E)**. Interestingly, the fraction of charged residues (FCR) also scales linearly with the isoform length, but not sequence distribution parameter, κ **(Fig S3E)**, which describes the concentration of similarly charged amino acids^43^ and was previously demonstrated as a valuable critical parameter to describe disordered protein’s statistical structures^44,45^. Nonetheless, a-Syn-115 has the lowest κ value of all 8 proteins, which indicates that similarly charged amino acids are more grouped than oppositely charged ones which may explain the structural predilection towards folding over itself more than the others **(Fig. 4A)**.

### Mapping the major cell types expressing *SNCA* transcript isoforms

Next, we mapped the various *SNCA* transcript isoforms to the major cell type/s in which they are expressed within each pathology group. For this analysis, we integrated the targeted MAS-Iso-Seq long read and unbiased snRNA-seq short read datasets. During cDNA generation, molecular identification barcode sequences specific to individual cells were integrated into each cDNA molecule. Thus, barcode sequences in MAS-Iso-Seq data could be matched to those from the same donor samples in the cell type-annotated snRNA-seq data. This allowed the mapping of the transcript isoform data to specific major cell types, including excitatory neurons, inhibitory neurons, astrocytes, microglia, oligodendrocytes, and OPCs **(Fig. 5A)**. Long-read data from 28 transcript isoform classes detected in 134 individual NC cells, 53 PD cells, and 219 DLB cells were mapped to annotated cell types. The distribution of *SNCA* transcript isoforms across the major cell-types in each disease state is summarized in **Table S8**. The vast majority of the long-read *SNCA* transcript isoforms were expressed in excitatory neurons. These results were consistent with the short-read snRNA-seq analysis described above, that demonstrated higher proportion of excitatory neurons expressing *SNCA* compared to other cell types **(Fig. 1B)**.

**Figure 5.**
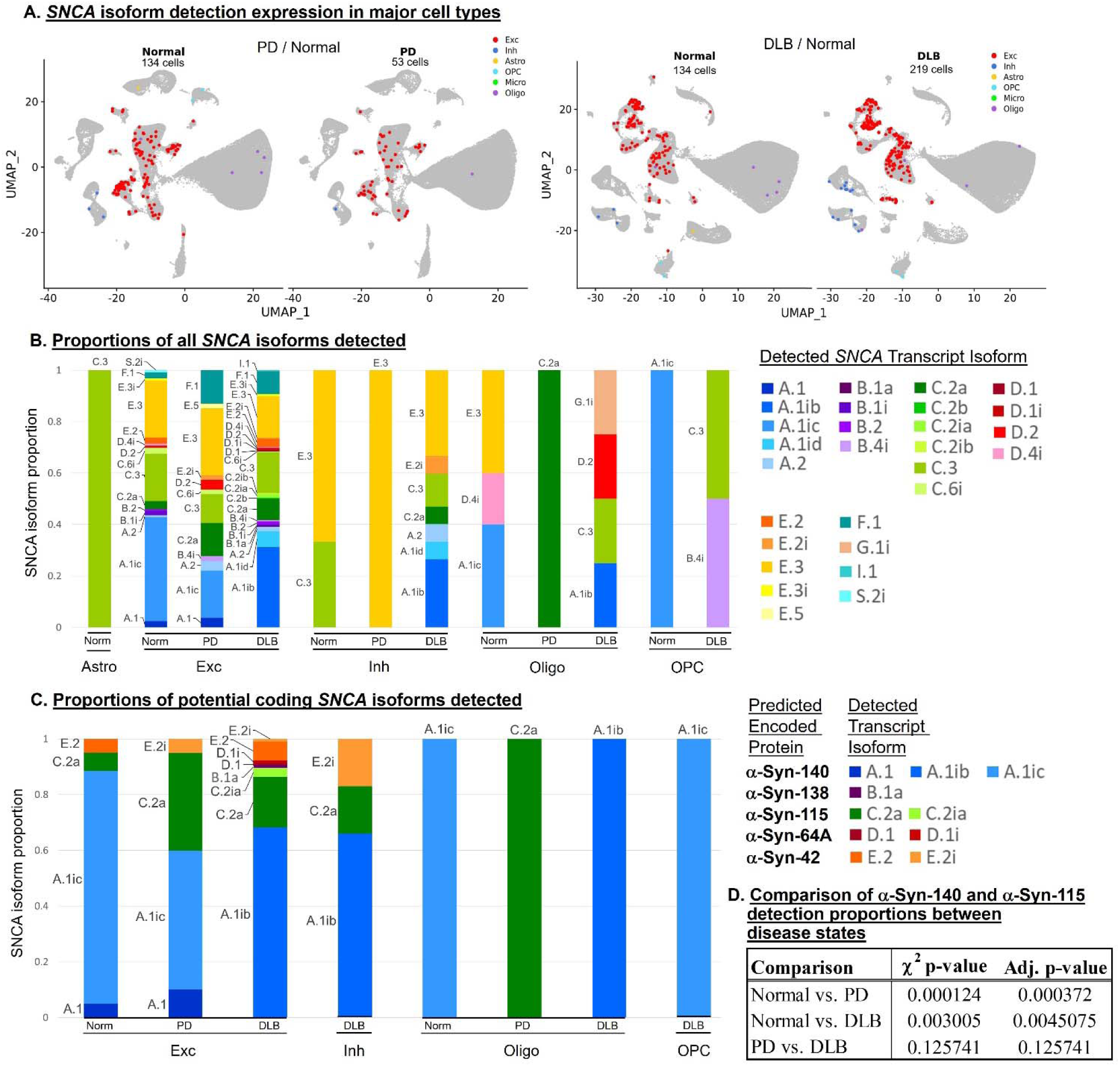
Detection of long-read *SNCA* transcript isoforms in major cell types. **A.** UMAP dimensional reduction plots of snRNA-seq transcriptomic data for PD (left panel) and DLB (right panel) nuclei integrated with NC nuclei. Plots are color coded to indicate nuclei with MAS-Seq long read data annotated as astrocytes (Astro), excitatory neurons (Exc), inhibitory neurons (Inh), microglia (Micro), oligodendrocytes (Oligo) and oligodendrocyte precursor cells (OPC). **B.** Histogram showing proportions of total transcript counts for transcripts of each isoform category of each major cell type and disease state with isoform detection. Color coding of isoform categories is shown to the right. **C.** Histogram showing proportions of total transcript counts for transcripts of each isoform category predicted to encode functional a-Syn protein (with exon 2-encoded membrane binding domain) of each major cell type and disease state with isoform detection. Color coding of isoform categories predicted to encode indicated protein variants is shown to the right. **D.** Table showing statistical summary for Chi-squared comparisons of a-Syn-140-and a-Syn-115-encoding isoform count proportions within NC and PD, NC and DLB, and PD and DLB excitatory neuron nuclei. Chi-squared p-values and p-values adjusted for multiple testing via Bonferroni correction are shown.

Accordingly, excitatory neurons exhibited the highest number of different isoforms, including isoforms from 9 different classes, within which A, C, and E were the most prominent **(Fig. 5B**, **Table S8)**. In further analysis, we examined the cell type distribution of *SNCA* transcript isoforms that are likely to encode the potentially functional proteins α-Syn-140, -138, -120, -115, -64A, -64B, -48, and -42 discussed above. Of these, isoforms encoding α-Syn-140, -138, 115, 64A, and 42 could be mapped to cell types **(Fig. 5C)**. As expected, the highest number of protein-encoding isoforms were detected in excitatory neurons. Comparison across disease states revealed that in NC excitatory neurons, approximately 90% of transcripts detected were predicted to encode α-Syn-140, while less than 10% encoded α-Syn-115. In notable contrast, in PD excitatory neurons, only 60% of transcripts encoded α-Syn-140, while 35% encoded α-Syn-115. Similarly, in DLB excitatory neurons, 70% encoded α-Syn-140, with 20% encoding α-Syn-115. Comparisons of proportional detection levels of isoforms predicted to encode α-Syn-140 and α-Syn-115 between NC and PD excitatory neurons, and between NC and DLB excitatory neurons, indicated that both PD and DLB excitatory neurons showed significantly higher proportions (*p_ad_*_j_ < 0.01) of α-Syn-115-encoding isoforms compared to NC **(Fig. 5D)**. The same comparison between PD and DLB excitatory neurons showed no significant difference (*p_ad_*_j_ > 0.1). Isoforms encoding α-Syn-42 were also detected in lower proportion in excitatory neurons across all disease states, while α-Syn-138 and α-Syn-64A-encoding isoforms were only detected in DLB excitatory neurons **(Fig. 5C)**. Inhibitory neurons from DLB also expressed isoforms encoding α-Syn-140, -115 and -42, but no functional α-Syn-encoding isoforms were detected in NC or PD inhibitory neurons. Only a maximum of 1 α-Syn-encoding isoform class was detected in each of the other cell types per disease state.

### Characterization of the excitatory neuronal subtypes expressing the diverse *SNCA* transcript isoforms

As described above, *SNCA* transcript isoforms were the most abundant in excitatory neurons. Thus, we sought to examine isoform expression more granularly within specific excitatory neuron subtypes. A total of 125 NC, 51 PD, and 200 DLB excitatory neurons with MAS-Iso-seq data were mapped via barcode matching to 14 subtype clusters derived from snRNA-seq data **(Fig. 6A)**. The largest numbers of cells expressing *SNCA* transcript isoforms were the Pyramidal Cell 1, Glutamatergic Neuron 1, 2, and 3, and Cajal-Retzius Cell 1 subtypes, with only a few cells expressing *SNCA* in the remaining clusters. Transcript isoform classes A, C, and E were the most abundant across all excitatory subtypes **(Fig. 6B**, **Table S9)**. Other isoforms were detected in several subtypes in either all or particular disease groups.

**Figure 6.**
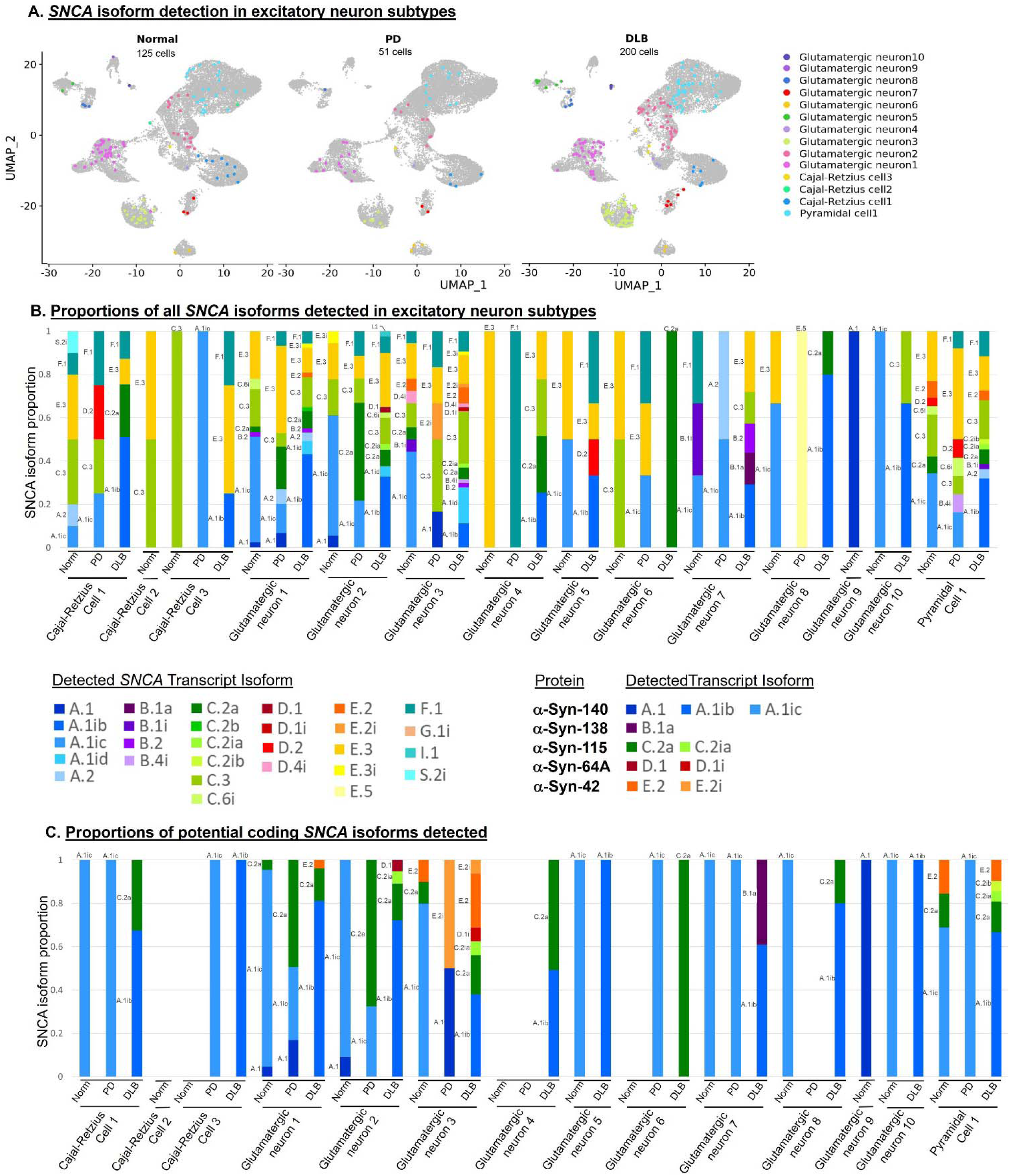
Detection of long-read *SNCA* transcript isoforms in excitatory neuron subtypes. **A.** UMAP dimensional reduction plots of snRNA-seq transcriptomic data for PD (left panel) and DLB (right panel) nuclei integrated with NC nuclei. Plots are color coded to indicate nuclei with MAS-Seq long read data annotated as indicated excitatory neuron subtypes. **B.** Histogram showing proportions of total transcript counts for transcripts of each isoform category of each excitatory neuron subtype and disease state with isoform detection. Color coding of isoform categories is shown below. **C.** Histogram showing proportions of total transcript counts for transcripts of each isoform category predicted to encode functional α-Syn protein (with exon 2-encoded membrane binding domain) of each excitatory neuron subtype and disease state with isoform detection. Color coding of isoform categories predicted to encode indicated protein variants is shown above.

Analysis in the context of functional proteins (as discussed above) showed that isoforms encoding a-Syn-140 were expressed at the highest proportion in the majority of excitatory neuron subtypes. However, isoforms encoding α-Syn-115 were predominant within specific subtypes and disease states, including PD Glutamatergic Neuron 1 and 2 cells, and DLB Glutamatergic Neuron 4 and 6 cells **(Fig. 6C**, **Table S9)**. However, in the corresponding NC subtypes, α-Syn-115-encoding transcripts were either not detected or were detected only in very low proportions. Similarly, α-Syn-42-encoding transcripts were prevalent in the Glutamatergic Neuron 3 subtype of both PD and DLB, but were rare in NC of this neuronal subtype. Isoforms encoding α-Syn-138 were only detected in DLB cells of the Glutamatergic Neuron 7 subtype, and isoforms encoding α-Syn-64A were only detected in DLB cells of the Glutamatergic Neuron 2 subtype.

## DISCUSSION

In this study, for the first time, we used a combined approach of targeted long-read and unbiased short-read single-cell sequencing to obtain new insights into the molecular mechanisms underlying α-Syn pathology across multiple synucleinopathies. These insights included 1) characterization of the full *SNCA* transcript isoform landscape in PD, DLB, and normal controls, 2) identification of predicted α-Syn protein variants expressed and their relative proportions across disease groups, 3) modeling of structural and biochemical properties of α-Syn protein variants to predict specific roles in pathogenesis, and 4) mapping of transcript and protein isoform expression to specific cell types and subtypes.

Through the combination of snRNA-seq and *SNCA*-targeted MAS-Iso-seq, we were able to identify a total of 327 unique *SNCA* transcript isoforms, compared to 41 isoforms in our previous bulk sequencing study^7^. This allowed us to more completely characterize the full landscape of *SNCA* isoforms, including rare transcripts. We thus identified 62 distinct splicing variant classes, the majority of which had not been previously reported, to our knowledge. These classes incorporated novel combinations of canonical exons, as well as noncanonical exons. In addition to 10 noncanonical exons identified within the intron 4 region and 2 downstream of exon 6, we also identified 2 novel exons located within the intron 3 region, both included within a single transcript. Characterizing this wider array of unique transcript isoforms also enabled us to predict novel α-Syn protein variants encoded by specific isoform classes. This led to the identification of 8 distinct α-Syn protein variants with potentially important mechanistic implications for disease. As noted above, we did not identify transcripts predicted to encode previously reported protein variants α-Syn-126, α-Syn-112, or α-Syn-98. However, we did identify one isoform, that may represent an incomplete α-Syn-112-encoding transcript. This isoform was highly expressed in one DLB sample, lending support to the possibility that it may be a significant contributor to α-Syn production in some circumstances. In our previous bulk tissue MAS-Iso-seq study^7^, transcripts encoding α-Syn-126 and α-Syn-98 were also not reported and, unlike α-Syn-112, these proteins have not been detected *in vivo*, suggesting that they may be relatively rare splicing variants.

The identification of α-Syn-115 was of particular interest due to both its predicted biochemical properties and its pattern of distribution across different disease states and among cell types and subtypes. The fact that transcript isoforms predicted to encode α-Syn-115 were detected across the widest range of donor samples of any α-Syn-encoding transcript class, including those encoding canonical α-Syn-140, suggests that this is not a very rare transcript and instead is consistently expressed, albeit likely at a significantly lower level than α-Syn-140. That multiple transcripts predicted to encode α-Syn-115 were also identified in our previous bulk tissue study^7^ lends further support to this possibility. It is likewise noteworthy that a greater proportion of *SNCA* transcripts in excitatory neurons appeared to encode α-Syn-115 relative to α-Syn-140 in both PD and DLB tissues compared to controls, suggesting that increased abundance of α-Syn-115 could contribute to synucleinopathy in these diseases. This potential is supported by our *in silico* predictions of structural and biochemical properties of α-Syn-115, wherein this protein variant is expected to display greater self-interaction affinity and greater affinity for interaction with α-Syn-140, suggesting the potential for increased cytosolic α-Syn-115 abundance to induce aggregation between nearby heterogeneous α-Syn isoforms. This also aligns with empirical findings for similarly-structured proteins, wherein other α-Syn variants lacking protective C-terminal disordered domains encoded by exons 5 and 6, and retaining aggregative central domains encoded by exons 3 and 4, such as α-Syn-112^23^ and the caspase-cleaved N103 protein^46^, have been shown to be more prone to fibril formation, as well as to induce parkinsonism in mice. Further work is needed to confirm the presence of α-Syn-115 in human tissues *in vivo* and to validate its predicted structural and biochemical properties through *in vitro* expression and analysis.

In addition to identifying increased proportional abundance for α-Syn-115-encoding isoforms in PD and DLB excitatory neurons overall, we were also able to pinpoint specific excitatory neuron subtypes that showed especially high proportional expression of α-Syn-115, including several glutamatergic neuron subtypes for which α-Syn-115-encoding transcripts showed equal or even greater detection levels than for α-Syn-140 in PD and DLB samples. It is also notable that while overall detection of α-Syn-42-encoding transcripts was low, in a particular glutamatergic neuron subtype, abundance of this transcript class was similar to that of α-Syn-140 in both PD and DLB samples, but qualitatively lower in control samples. Another subtype showed proportionally high expression of α-Syn-64A-encoding transcripts in DLB. Thus, we observe that expression of atypical, potentially aggregative α-Syn-encoding isoforms does not appear to be evenly distributed across excitatory neuron subtypes, but rather that specific subtypes show especially high levels of these isoforms, suggesting that these subtypes may represent focal points for fibril formation and drivers of disease.

The findings of this study have translational implications for the development of new precision medicine strategies to combat synucleinopathies, indicating the potential for treatments targeting both specific transcript and protein isoforms, as well as disease-driving cell subtypes. Treatments employing antisense oligonucleotides and small molecules to modify or degrade specific mRNA species have been previously developed for treatment of spinal muscular atrophy^47,48^, ALS^49^, muscular dystrophy^50^, and cancer^51^, while RNA interference has been used to degrade harmful mRNA in treatment of amyloidosis^52^, among other diseases^53,54^, and CRISPR-Cas13 systems have also been employed for knockdown of specific RNA targets^55^. It may be possible to adapt these or other methods to alter or degrade aberrantly spliced *SNCA* transcript isoforms identified here to help prevent or control the progression of synucleinopathy by reducing accumulation of aggregative α-Syn protein variants. Additionally, these protein variants such as α-Syn-115 could be directly targeted for proteolysis or to prevent aggregation using antibody-based methods as in AD^56^ and breast cancer^57^ treatments, proteolysis targeting chimeras as employed in prostate cancer models^58^, or other methods. Further characterization of disease-driving cell subtypes identified in this study could also enable the development of treatment strategies targeting these specific subtypes, as has been studied for cell types such as disease-associated microglia in AD^59–61^. The detection of high proportions of alternatively spliced transcripts in PD and DLB samples also opens up the possibility that the presence of alternatively spliced *SNCA* transcripts or protein variants in cerebrospinal fluid^62^ or in the gut^63^ could potentially serve as biomarkers for early detection of synucleinopathies prior to disease onset and thereby facilitate treatment strategies aimed at preventing or reducing the impact of disease.

This work provides an important investigation into the transcriptional variation of *SNCA* across multiple synucleinopathies. However, there are some limitations. First, as noted above, our cDNA generation method using polyA targeting, while highly effective in capturing mRNA, is intrinsically 3’ biased, potentially leading to molecules with incomplete 5’ ends. As described above, we have addressed this issue by treating isoforms lacking 5’ UTRs as potentially incomplete and grouped incomplete transcripts with matching complete isoform categories. Second, because we used a targeted MAS-Iso-seq approach to enrich our samples for *SNCA* and maximize detection of *SNCA* transcript isoforms, accurate quantification of isoform expression levels is difficult, as our enrichment process may not uniformly impact all variant transcripts. Furthermore, the aforementioned potential for incomplete cDNA generation may also result in increased abundance for shorter transcripts compared to longer ones. For these reasons, we do not attempt to precisely quantify isoform expression levels here. However, detection of a specific isoform class across numerous donor samples is highly suggestive of consistently robust expression. Furthermore, as we should be able to assume that the relative capacity for detection of specific sets of isoforms (such as those encoding α-Syn-140 and α-Syn-115) remains approximately equivalent across samples, it is possible to make valid comparisons of the detection ratios of these isoform sets between samples.

Understanding the precise molecular mechanisms through which *SNCA* contributes to the onset and progression of synucleinopathies such as PD and DLB in specific cell types is crucial. In this manuscript, we have characterized the *SNCA* transcriptomic landscape across multiple synucleinopathies at an unprecedented cell subtype level of precision, and presented evidence that expression of alternatively spliced *SNCA* isoforms may be proportionally increased in the context of disease, potentially leading to higher proportions of protein variants with increased aggregation propensity. This work demonstrates the utility of combining short and long-read single-cell sequencing methods to catalogue transcript isoforms within specific cell types and subtypes and to compare these across disease states. However, examination of *SNCA* alone does not tell the full story of the molecular genetic mechanisms underlying neurodegeneration in synucleinopathies. It will be important in future work to expand on these findings by applying this methodology to compare the whole transcriptomic full-length isoform landscapes across disease states and within specific cell subtypes in order to more thoroughly understand the posttranscriptional mechanisms driving pathology and thereby facilitate the development of effective strategies for prevention and treatment of these devastating diseases.

## METHODS

### Human post-mortem brain tissue samples

The demographics, pathological notes, and other metadata for this study cohort are detailed in **Table S1**. Extended pathology information for PD samples is provided in **Table S2**. Frozen human TC tissue samples from donors clinically diagnosed with DLB (*n* = 12) and NC donor samples (*n* = 12) were obtained from the Kathleen Price Bryan Brain Bank (KPBBB) at Duke University. Samples from donors diagnosed with PD (*n* = 13) were obtained from the Banner Sun Health Research Institute (BSHRI)^64^. NC samples were derived from donors with no clinical history of neurological disorder and samples had no neuropathological evidence of neurodegenerative diseases. All DLB donors were confirmed to exhibit Lewy-related pathology within the neocortical, limbic, or brainstem regions and showed low levels of AD neuropathologic change (Braak stages I or II), with the exception of donor 1097 which exhibited Braak stage III pathology. Donor patient PD diagnoses were defined by the presence of two of the three cardinal clinical signs of resting tremor, muscular rigidity and bradykinesia. Additionally, diagnoses of all PD samples were confirmed in autopsy by observation of pigmented neuron loss and the presence of Lewy bodies in the substantia nigra (SN). Neuropathological states of PD samples were confirmed *postmortem* using established clinical practice recommendations for McKeith scoring^65^ and staging via the Unified Staging System for Lewy Body Disorders (USSLB)^66^. All PD samples for which information was available had McKeith scores ranging from moderate to severe (2-4) in both the amygdala and SN. Where available, TC McKeith scores for most of the PD samples were either 0-1, with one sample each receiving scores of 2 and 3, indicating mild or absent PD pathology in this region for the majority of samples. USSLB stages of PD samples ranged from II-IV. PD samples 96-36 and 96-49 were lacking specific USSLB stage determination due to harvesting prior to BSHRI standardization of stage determination protocol. All tissue donors were Caucasians with the *APOE* e3/e3 genotype. The project was approved for exemption by the Duke University Health System Institutional Review Board. The methods described were conducted in accordance with the relevant guidelines and regulations.

### Nuclei isolation from post-mortem human brain tissue

The nuclei isolation procedure has been described^67^, and was based on previous studies^68,69^ and optimized for single-cell experiments. 100-200 mg of human TC brain tissue samples were thawed in Lysis Buffer (0.32 M Sucrose, 5 mM CaCl_2_, 3 mM Magnesium Acetate, 0.1 mM EDTA, 10 mM Tris-HCl pH 8, 1 mM DTT, 0.1% Triton X-100) and homogenized with a 7 ml dounce tissue homogenizer (Corning) and filtered through a 100 μm cell strainer, transferred to a 14 x 89 mm polypropylene ultracentrifuge tube, and underlain with sucrose solution (1.8 M Sucrose, 3 mM Magnesium Acetate, 1 mM DTT, 10 mM Tris-HCl, pH 8). Nuclei were separated by ultracentrifugation for 15 minutes at 4°C at 107,000 RCF. Supernatant was aspirated, and nuclei were washed with 1 ml Nuclei Wash Buffer (10 mM Tris-HCl pH 8, 10 mM NaCl, 3 mM MgCl_2_, 0.1% Tween-20, 1% BSA, 0.2 U/μL RNase Inhibitor). Resuspended nuclei were centrifuged at 300 RCF for 5 minutes at 4°C, and supernatant was aspirated. Nuclei were then resuspended in Wash and Resuspension Buffer (1X PBS, 1% BSA, 0.2 U/μL RNase Inhibitor), then filtered through a 35 μm strainer. Nuclei concentrations were determined using a Countess™ II Automated Cell Counter (ThermoFisher) and nuclei quality was assessed at 10X and 40X magnification using an Evos XL Core Cell Imager (ThermoFisher).

### cDNA library preparation and snRNA-seq

snRNA-seq libraries were constructed as previously^67^ using the Chromium Next GEM Single Cell 3’ GEM, Library, and Gel Bead v3.1 kit, Chip G Single Cell kit, and i7 Multiplex kit (10X Genomics) according to manufacturer’s instructions. For each sample, 10,000 nuclei were targeted. Library quality control was performed on a Bioanalyzer (Agilent) with the High Sensitivity DNA Kit (Agilent) according to manufacturer’s instructions and the 10X Genomics protocols. Libraries were submitted to the Sequencing and Genomic Technologies Shared Resource at Duke University for quantification using the KAPA Library Quantification Kit for Illumina® Platforms and sequencing. Groups of four snRNA-seq libraries were pooled on a NovaSeq 6000 S1 50bp PE full flow cell to target a sequencing depth of 400 million reads per sample (Read 1 = 28, i7 index = 8, and Read 2 = 91 cycles). Sequencing was performed blinded to age, sex, and diagnosis.

### snRNA-seq data processing

Raw snRNA-seq sequencing data were converted to FastQ format, aligned to a GRCh38 pre-mRNA reference, filtered, and counted using CellRanger 4.0.0 (10X Genomics). Subsequent processing was done using Seurat 4.0.1^70^. Filtered feature-barcode matrices were used to generate Seurat objects for the individual samples. For QC filtering, nuclei below the 1^st^ and above the 99^th^ percentile for number of features were excluded. Nuclei above the 95^th^ percentile for mitochondrial gene transcript proportion (or >5% mitochondrial transcripts if 95^th^ percentile mitochondrial transcript proportion was <5%) were also excluded. Because experiments were conducted in nuclei rather than whole cells, mitochondrial genes were subsequently removed. The individual sample Seurat objects were merged into one, and were iteratively normalized using SCTransform^71^ with glmGamPoi, which alleviates bias from weakly-expressed genes^72^. Batch correction was performed using reference-based integration^35^ on the individual sample normalized datasets, which improves computational efficiency for integration.

### Doublet/Multiplet detection in snRNA-seq data

Multiplets comprising different cell types (heterotypic) were excluded from snRNA-seq data by considering the “hybrid score”, as described previously^67^. The hybrid score is calculated as (x_1_ – x_2_) / x_1_, where x_1_ is the highest and x_2_ is the second highest prediction score^73^. Heterotypic multiplets would be expected to exhibit competing cell type prediction scores due to the presence of transcriptomic/epigenomic profiles from multiple cell types. Multiplets composed of one cell type (homotypic) were identified based on the number of features per cell. snRNA nuclei with feature counts > 99^th^ percentile were excluded. Removal of homotypic multiplets in this manner is expected to also aid in filtering of heterotypic multiplets.

### Cell type and subtype cluster annotation

Cell type annotation was conducted using a label transfer method^35^ and a previously annotated reference dataset from human M1. Batch-corrected data from both our dataset and the human M1 dataset were used for label transfer. Nuclei with maximum prediction scores of <0.5 were excluded. Nuclei with a percent difference of <20% between first and second highest cell type prediction scores were termed “hybrid” and excluded^73^. Endothelial cells and vascular leptomeningeal cells (VLMCs) were in low abundance and did not form distinct uniform manifold approximation and projection (UMAP) clusters and were thus excluded. Proportion values of each cell type were tested for normal distribution using the Shapiro-Wilk normality test and found likely to be normally distributed. One-way analysis of variance (ANOVA) was used to test for group differences in mean cell type proportions between diagnosis groups for each major cell type. Two-tailed t-tests were used to compare differences in mean cell type proportions between samples obtained from the Banner Sun Health Research Institute Brain and Body Donation Program (PD samples) and the Duke Alzheimer’s Disease Research Center (ADRC) Kathleen Price Bryan Brain Bank (NC, AD, and DLB samples). ANOVA and t-test p-values for the six major cell types were then adjusted for multiple testing using the Benjamini-Hochberg false discovery rate (FDR) method. Following principal component analysis (PCA), dimensionality was examined using an Elbow plot and by calculating variance contribution of each PC. UMAP was then run using the first 30 PCs, and nuclei were clustered based on UMAP reduction. The resolution levels for cluster delineation were selected after comparison of a range of values as it was determined to provide optimal distinction between populations of nuclei displaying unique gene expression profiles as evidenced by their separation from one another in UMAP space. Counts of predicted major cell types based on the label transfer were examined for each of the clusters, and clusters were manually annotated based on the majority cell type for each cluster (e.g., ‘Exc1’, ‘Exc2’, etc.). Proportions of nuclei belonging to each cell type were calculated for each donor sample by dividing the number nuclei annotated with a particular cell type by the total number of nuclei from that sample in the dataset. Where noted, clusters were also subsequently annotated using the scMayoMap^37^ R package.

### Human M1 reference data processing

To optimize label transfer, we re-processed previously published human primary motor cortex (M1) snRNA-seq data^36^ to map it to GRCh38 Ensembl 80 as we did with our data^67^. FastQ files were obtained from the Neuroscience Multi-omic Data Archive (NeMO: https://nemoarchive.org/) and were aligned to the same GRCh38 pre-mRNA reference used for our data, filtered, and counted using CellRanger 4.0.0 (10X Genomics). Filtered feature-barcode matrices were used to generate separate Seurat objects for each sample, with nuclei absent from the annotated metadata excluded. Seurat objects were merged and iteratively normalized using SCTransform^71^ with glmGamPoi. Batch correction was performed using reference-based integration^35^ on the normalized datasets. The 127 transcriptomic cell types in this data were grouped into 8 broad cell types including astrocytes, endothelial cells, excitatory neurons, inhibitory neurons, microglia, oligodendrocytes, OPCs, and VLMCs.

### Targeted Mas-Seq sequencing

The Mas-Seq libraries were prepared from the 12 cDNA libraries of Chromium Next Gem Single Cell 3’ kit. Custom probes and blockers were designed using Twist® hybridization capture for the gene *SNCA*. TSO artifacts were removed using PacBio ® MAS Capture primers and MAS capture beads. The capture beads/DNA complex was used as input in the Twist® pre-capture amplification PCR, using custom primers. Twist® post-capture amplification was performed using the same custom PCR primers used in the pre-capture reaction. The post-capture amplified, target enriched samples then underwent PacBio® MAS PCR, MAS Array Formation, DNA damage repair, and nuclease treatment. The samples were quantified using Qubit® 3.0 after each step and analyzed using TapeStation (Agilent®) after pre-capture amplification, post-capture amplification, MAS PCR, MAS Array formation to track performance throughout the process. The target-enriched MAS-Seq libraries were loaded into one SMRTcell-8M each at 150-200pM on-plate loading concentration and run on the Sequel-IIe in MAS-Seq Single Cell mode with movie duration of 30 h in length. In two cases, libraries were sequenced on the Revio for comparison, in one SMRT-Cell-25M each at 250pm on-plate loading concentration with movie duration of 30 h in length. All sequencing was performed at Eremid Genomic Services.

### MAS-Iso-Seq data processing

Raw MAS-Seq data was processed on SMRT ® Link v11.0 software^74^ using the Iso-Seq single-cell command line interface (CLI) workflow (https://isoseq.how/umi/cli-workflow.html). The software segmented the HiFi reads and extracted primers, unique molecular identifiers (UMIs), and barcode information. The barcodes were corrected for barcode errors with single cell whitelists (10x Genomics 3’kit v3.1). The UMIs were deduplicated for PCR duplication by clustering for UMI and cell barcode. Redundant isoforms collapsed and unique isoforms were annotated. Further processing was performed using the Pigeon PacBio Transcript Toolkit software and classification CLI workflow (https://isoseq.how/classification/workflow.html). The reads were mapped to chromosomal regions by aligning to reference genome GRCh38. The transcripts were then classified via SQANTI3 categories. The isoforms were then filtered for primer annealing to downstream A-rich regions (IntraPriming) and artifacts of reverse transcriptase template switching (RTSwitching) and converted into a gene and isoform count matrix to create a Seurat Object per sample. Seurat 5.0.0 and version 4.1.0 were used for further processing. All 12 samples were merged based on similar exon structures. The coordinates of the exons were used from the gff generated from collapsed isoforms.

### Transcript isoform classification

Isoforms were manually grouped based on general splicing attributes, including the specific exons included in the transcripts, as well as whether the splice sites between exons were canonical or extended or truncated at the 5’ or 3’ ends. The inclusion of predicted ORFs within transcripts was also taken into consideration. Each isoform class was assigned a letter identification based on the set of exons included or potentially included in the transcript, and each unique splice pattern within this exon-set was assigned a number. Because cDNA molecules in this study were generated from mRNA by 3’ to 5’ extension via reverse transcription from polyA tail-annealing primers, the long-read sequences subsequently generated display a 3’ bias and may thus have incomplete 5’ ends. For this reason, transcripts were considered to be potentially incomplete unless they included 5’ UTR upstream of the canonical ATG translation start codon in exon 2. Potentially incomplete transcripts were designated with a lowercase letter “i” in their transcript IDs and were grouped into classes as if they included all canonical exons upstream of their 5’ termini and were assigned the same number as any complete transcript with the same splicing pattern as their sequenced region. In cases where multiple potentially incomplete transcripts could match to the same complete transcript, these were additionally distinguished by assignation of a lowercase italicized letter “*a*-*d*” in their IDs. Lowercase letters were also used to distinguish transcripts predicted to include ORFs encoding potentially functional α-Syn protein from those that were not, based on the presence of the structured N-terminal membrane-binding domain encoded on exon 2. ORFs were detected within each transcript using the ORFik R software package^75^. Major categories were assigned based on which exons were included in the transcript. Sub-categories were annotated based on splicing patterns of 5’ and 3’ ends of each exon, which could be canonical, extended, or truncated. Isoforms were categorized as complete if they contained 5’ UTR upstream of the canonical *SNCA* start codon within exon 2, and incomplete if they did not. Incomplete isoforms were grouped with complete isoforms with the same splicing patterns as those present in the incomplete isoform, with the exception of the 5’ end of the first exon present in the incomplete transcript, which needed to have the same splicing or be truncated compared to the same exon in the corresponding complete isoform category.

### Assessment of isoform abundance

Raw sequencing counts were log normalized **(Table S5)**. When assessing abundance of transcripts encoding potentially functional α-Syn protein, counts from all transcript classes predicted to encode the same protein variant were summed. Additionally, counts from incomplete isoform categories likely to encode the same protein were pooled with the corresponding complete isoform class counts. Incomplete isoforms classes were only considered likely to encode a particular α-Syn variant if they did not potentially match to any complete isoform classes other than the coding class, and also were not mono-exonic. This included A.1 and A.1i*a-c* for α-Syn-140, B.1 for α-Syn-138, P.1 for α-Syn-120, C.2*a*, C.2i*a*, L.1, and O.1 for α-Syn-115, D.1 and D.1i for α-Syn-64A, H.1 for α-Syn-64B, R.1 for α-Syn-48, and E.2 and E.2i for α-Syn-42.

### *In silico* structural modeling of predicted s-Syn protein variants

Transcript sequences were translated into amino acid sequences using the NCBI ORF Finder with default parameters. The resulting protein sequences were input into AlphaFold2 via ColabFold v1.5.5^76^ using the MMseqs2 pipeline with default settings for structure prediction. Images were generated from PDB outputs using The PyMOL Molecular Graphics System, Version 3.1, Schrödinger, LLC. Disorder scores were calculated using Metapredict^40^. Predicted intermolecular interaction maps (intermaps) were generated using the FINCHES online tool (https://finches-online.com/intermaps), applying the Mpipi-GG method with a window size of 9. The value of the mean-field intermolecular interaction parameter (ε) was calculated as described^42^.

### Statistical comparisons

Comparisons of proportions of cells expressing *SNCA* between major cell types - astrocytes, excitatory neurons, inhibitory neurons, microglia, oligodendrocytes, and oligodendrocyte precursor cells (OPCs) - were carried out by concatenating proportions for each cell type across all donor samples, then using one-way ANOVA tests with post-hoc Tukey HSD tests for paired comparisons between each pair of cell types. Comparisons of proportions of cells expressing *SNCA* between disease states (NC, PD, and DLB) within each major cell type were carried out by concatenating proportions for each cell type from all donor samples of a particular disease state, then using one-way ANOVA tests with post-hoc Tukey HSD tests for paired comparisons between each pair of disease states. Comparisons of the relative proportions of the detection levels of transcript isoforms predicted to encode α-Syn-140 and α-Syn-115 between NC and PD excitatory neurons, and between NC and DLB excitatory neurons were carried out using Chi-squared tests with Bonferroni correction for multiple testing.

### Genome version and coordinates

All genomic data and coordinates are based on the December 2013 version of the genome: hg38, GRCh38. Genomic DNA sequence from sample 96_36^7^ was converted from hg19 coordinates to hg38 via UCSC genome liftover^35^.

## DECLARATIONS

### Ethics approval and consent to participate

The project was approved by the Duke Institutional Review Board. The study does not involve living human subjects. All samples were obtained from autopsies, and all are de-identified.

### Consent for publication

Not applicable.

### Availability of data and materials

The snRNA-seq count data and MAS-Iso-seq isoform data generated in this study will be available at the Synapse data repository (https://synapse.org, ProjectSynID: syn50996869 and syn60245188). Access will be avaliable upon publication under controlled use conditions. All computer code used for this study will be available before publication on GitHub (https://github.com/chibafaleklab).

## Funding

This work was funded in part by the National Institutes of Health/National Institute of Neurological Disorders & Stroke (NIH/NINDS) [RF1-NS113548-01A1 to OC-F] and by the National Institutes of Health/National Institute on Aging (NIH/NIA) [R01 AG057522 and RF1 AG077695 to OC-F]. The Brain and Body Donation Program has been supported by the National Institute of Neurological Disorders and Stroke (U24 NS072026 National Brain and Tissue Resource for Parkinson’s Disease and Related Disorders), the National Institute on Aging (P30 AG019610 and P30AG072980, Arizona Alzheimer’s Disease Center), the Arizona Department of Health Services (contract 211002, Arizona Alzheimer’s Research Center), the Arizona Biomedical Research Commission (contracts 4001, 0011, 05-901 and 1001 to the Arizona Parkinson’s Disease Consortium) and the Michael J. Fox Foundation for Parkinson’s Research .

### Authors’ contributions

O.C-F. conceived of the presented idea. O.C-F. acquired brain samples and designed the study sample. G.E.S. and T.G.B. provided PD brain tissue and associated data. D.C.G. and J.G. performed tissue processing, nuclei extraction, and the 10X experiments for snRNA-seq library preps. E.K.S., W.P., and D.C.G. processed MAS-Iso-seq data. Z.M. processed snRNA-seq data. E.K.S., W.P., and D.C.G. performed MAS-Iso-seq data analyses. S.M., D.Y., A.S, Y.E., and R.B. performed protein structural modeling and predictive computational biochemical analyses. O.C-F., Y.E., and R.B. planned and supervised the work. E.K.S., W.P., and O.C-F. discussed and interpreted the results. E.K.S., W.P., S.M., D.Y., and A.S prepared figures and tables. E.K.S., W.P., O.C.-F, S.M., D.Y., A.S, Y.E., and R.B. wrote the manuscript. D.C.G. contributed to writing of the methods section. All authors read and approved the final manuscript. O.C-F. obtained funding support.

## Acknowledgements

We thank the Kathleen Price Bryan Brain Bank at Duke University (funded by NIH/NIA R01 AG028377) for providing us with the brain tissues, and the Duke Sequencing and Genomic Technologies Shared Resource for sequencing. We are grateful to the Banner Sun Health Research Institute Brain and Body Donation Program of Sun City, Arizona for additional provision of human biological materials. R. B. thanks the Israel Science Foundation (1454/20) for the support of this research.

## LIST OF ABBREVIATIONS

(α-Syn): Alpha-synuclein
(PD): Parkinson’s disease
(DLB): dementia with Lewy bodies
(MAS-Iso-seq): long-read multiplexed arrays isoform sequencing
(sn): single nucleus
(fPD): familial form of PD
(GWAS): genome-wide association studies
(TSS): transcriptional start site
(UTR): untranslated region
(AD): Alzheimer’s disease
(ALS): amyotrophic lateral sclerosis
(aa): amino acids
(NC): normal control
(BSHRI): Banner Sun Health Research Institute
(USSLB): Unified Staging System for Lewy Body Disorders
(VLMCs): vascular leptomeningeal cells
(UMAP): uniform manifold approximation and projection
(ANOVA): analysis of variance
(ADRC): Alzheimer’s Disease Research Center
(FDR): false discovery rate
(PCA): principal component analysis
(NeMO): Neuroscience Multi-omic Data Archive
(CLI): command line interface
(UMI): unique molecular identifier
(ε): mean-field intermolecular interaction parameter
(OPC): oligodendrocyte precursor cell
(TC): temporal cortex
(QC): quality control
(ORF): open reading frame
(pLDDT): per-residue model confidence
(IDP): intrinsically disordered protein
(FCR): fraction of charged residues

---

**Figure S1.**
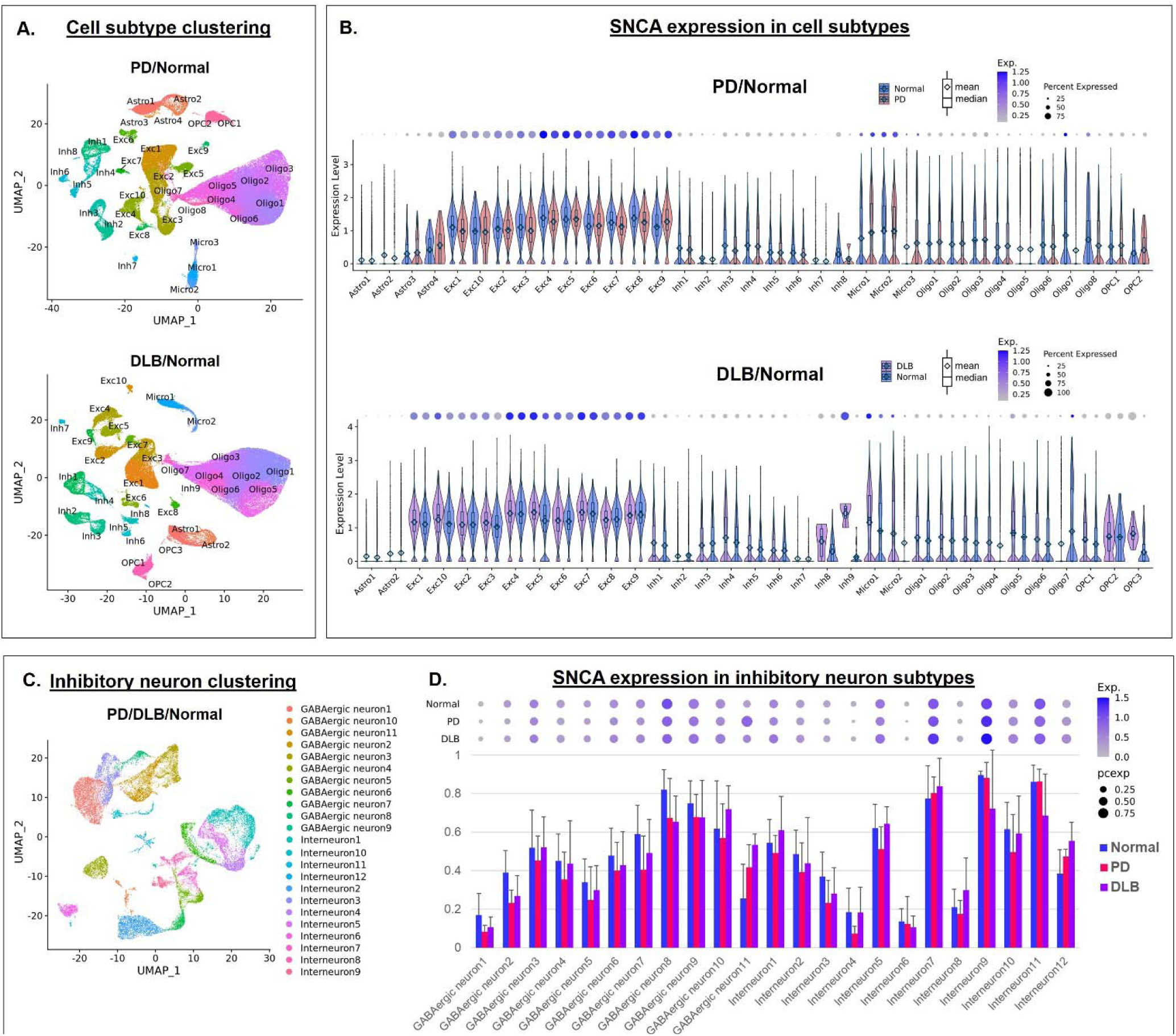
Patterns of SNCA expression in Normal, PD, and DLB excitatory and inhibitory neuron subtypes. **A.** UMAP dimensional reduction plots of snRNA-seq transcriptomic data for PD (upper panel) and DLB (lower panel) nuclei integrated with NC nuclei. Plots are color coded to indicate nuclei annotated as astrocyte (Astro), excitatory neuron (Exc), inhibitory neuron (Inh), microglial (Micro), oligodendrocyte (Oligo) and oligodendrocyte precursor cell (OPC) subtypes. **B.** Violin plots showing SNCA expression levels for individual nuclei within each disease state/cell subtype. Box plots represent upper and lower quartiles, with center lines indicating data median and diamonds indicating means and whiskers indicating data spread. **C.** UMAP dimensional reduction plot of snRNA-seq transcriptomic data for inhibitory neurons of PD and DLB nuclei integrated with NC nuclei. Plots are color coded to indicate subtype clusters with MayoMap annotation. **D.** Histogram showing proportions of nuclei of each inhibitory neuron subtype and disease state expressing SNCA. Bars represent mean proportions of expressing nuclei for each donor sample within corresponding disease state/cell type. Error bars represent standard deviations. Dot plots above histograms indicate SNCA expression level (color saturation) and proportion (dot size).

**Figure S2.**
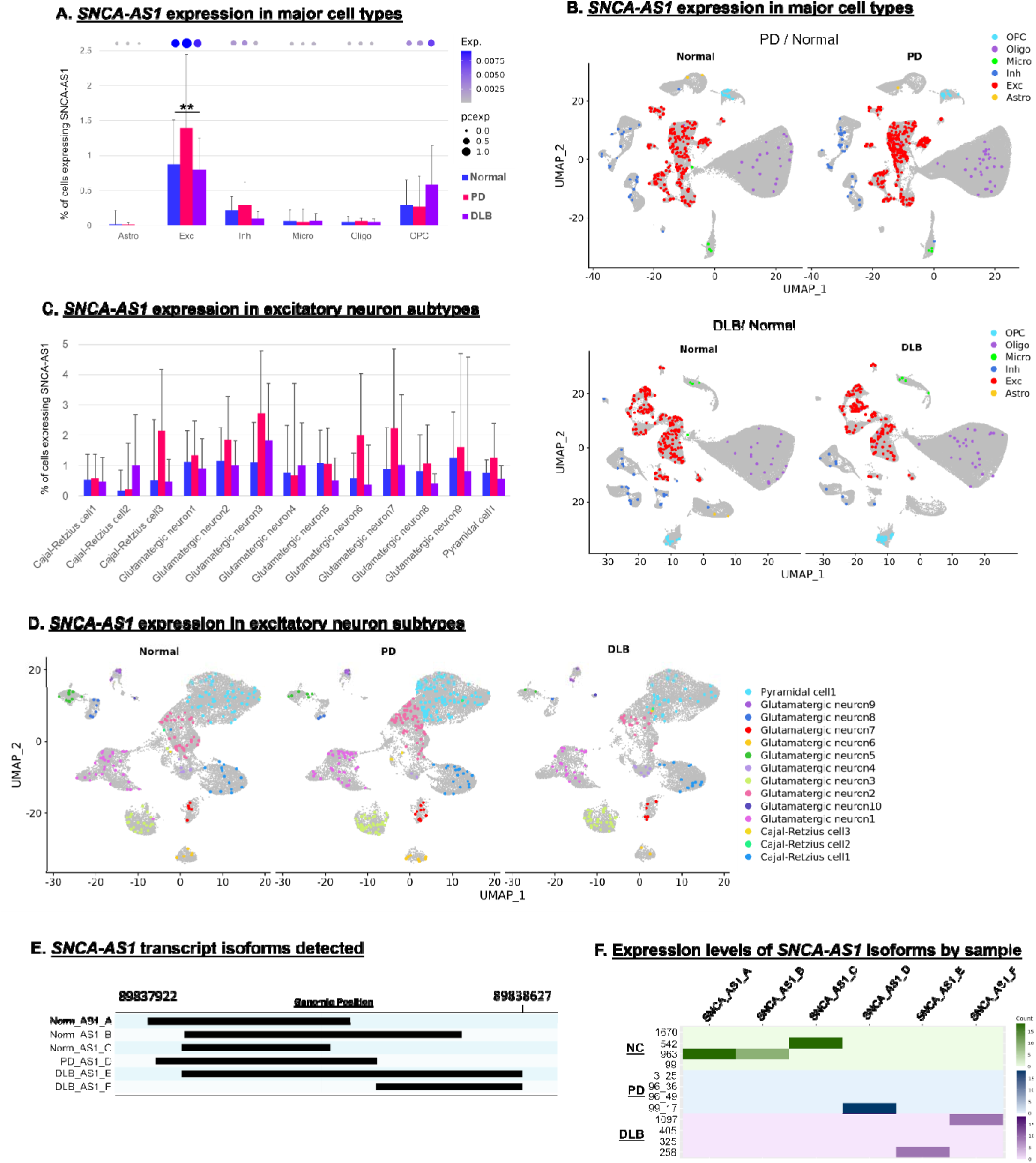
Expression patterns of *SNCA-AS1* in Normal, PD, and DLB cell types and subtypes. **A.** Histogram showing proportions of nuclei of each major cell type and disease state expressing *SNCA-AS1*. Bars represent mean proportions of expressing nuclei for each donor sample within corresponding disease state/cell type. Error bars represent standard deviations. **= p<0.01, Comparisons of pooled diagnosis groups between the 6 cell types via ANOVA with post-hoc Tukey tests and Bonferroni correction for multiple testing. Dot plots above histograms indicate *SNCA* expression level (color saturation) and proportion (dot size). **B.** UMAP dimensional reduction plots of snRNA-seq transcriptomic data for PD (upper panel) and DLB (lower panel) nuclei integrated with NC nuclei. Plots are color coded to indicate nuclei expressing *SNCA-AS1* annotated as astrocytes (Astro), excitatory neurons (Exc), inhibitory neurons (Inh), microglia (Micro), oligodendrocytes (Oligo) and oligodendrocyte precursor cells (OPC). **C.** Histogram showing proportions of nuclei of each excitatory neuron subtype and disease state expressing *SNCA-AS1*. Bars represent mean proportions of expressing nuclei for each donor sample within corresponding disease state/cell type. Error bars represent standard deviations. **D.** UMAP dimensional reduction plot of snRNA-seq transcriptomic data for excitatory neurons of PD and DLB nuclei integrated with NC nuclei. Plots are color coded to indicate nuclei expressing *SNCA-AS1* within subtype clusters with MayoMap annotation. **E.** Genomic track plots showing exon regions for *SNCA-AS1* transcript isoforms. Chromosome 4 region covered by track plots is indicate above plots. Alternating blue and light blue rows show unique isoform splicing categories, with identifiers shown to the left side. F. Heatmap showing log-normalized counts per donor sample of *SNCA-AS1* transcripts of each isoform panel E. NC samples are shown in shades of green, PD samples in shades of blue, and DLB samples in shades of purple.

**Figure S3.**
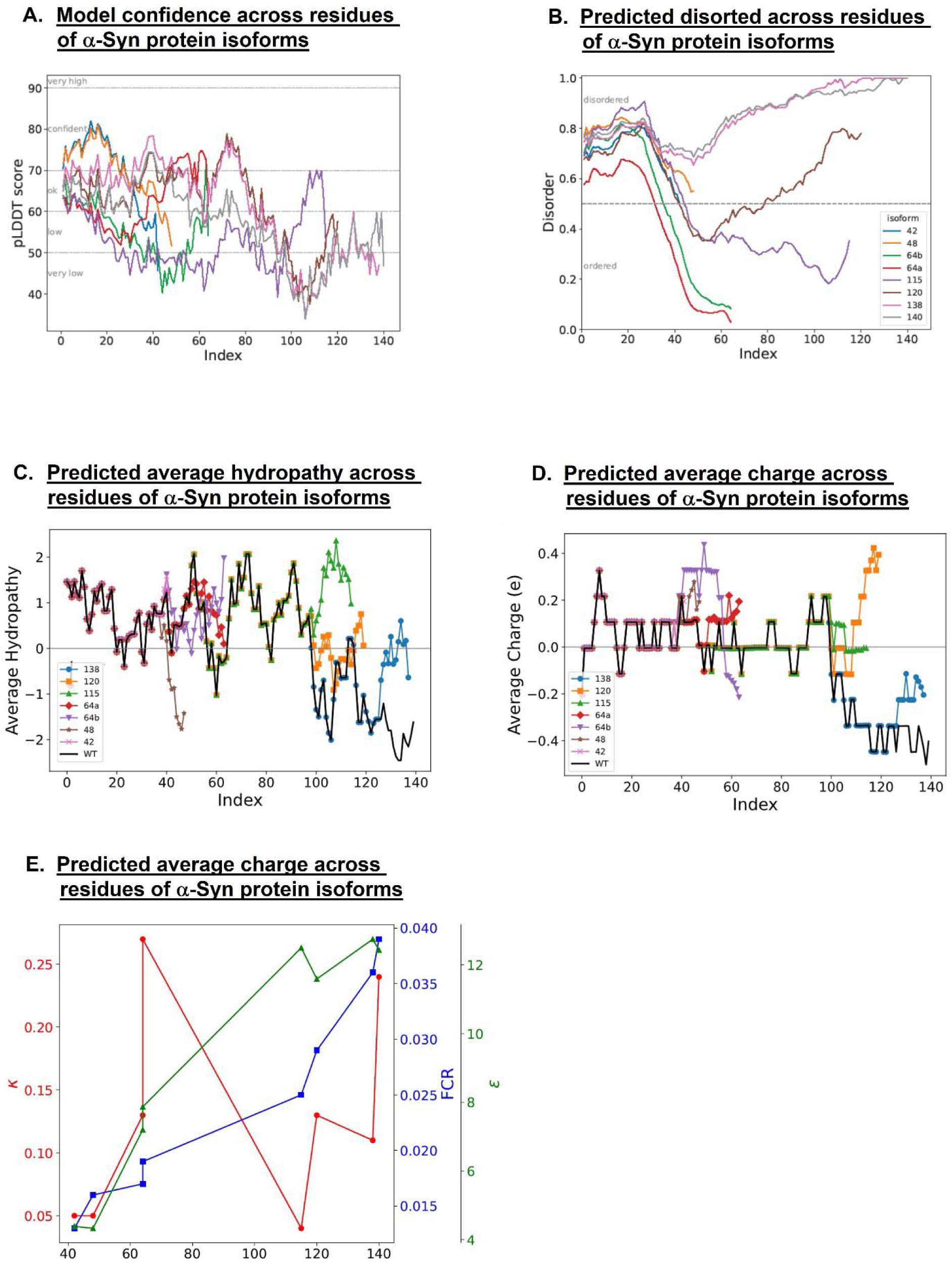
Structural modeling of predicted a-Syn protein variants. **A.** pLDDT confidence scores of the AlphaFold2 predicted structures. The N terminal is ordered and the C terminal is disordered. **B.** Predicted disorder scores for different unbound isoforms by Metapredict. The horizontal line defines the transition from an ordered section to a disordered one. In all cases, in the unbound state, the N terminal is predicted to be disordered in solution. **C.** Moving average hydropathy distribution calculated at pH 7 and room temperature. All isoforms longer than 100 amino acids are more hydrophobic than the canonic 140 long isoform. Moving average is over 5 amino-acids. WT = a-Syn-140. **D.** Moving average charge distribution calculated at pH 7 and room temperature. All isoforms have more positive charge distributions than a-Syn-140. WT = a-Syn-140. **E.** Sequence-based interaction and electrostatic characterization of the isoforms. Sequence distribution parameter, κ (red), fraction of charged residues, FCR (blue), and self-interaction estimation, ε (green) are shown for each isoform. With an increase of FCR and ε according to isoform’s length, there seems to be little correlation between the isoform length and κ value, with a-Syn-115 having the lowest value and the two pairs of similar length isoforms (64, 138-140) with significant different κ values.

